# Localization of T cell clonotypes using spatial transcriptomics

**DOI:** 10.1101/2021.08.03.454999

**Authors:** William H. Hudson, Lisa J. Sudmeier

## Abstract

Spatial transcriptomics is an emerging technology that measures gene expression while preserving spatial information. Here, we present a method to determine localization of specific T cell clones by obtaining T cell receptor (TCR) sequences from spatial transcriptomics assays. Our method uses an existing commercial spatial transcriptomics platform and open-source software for analysis, allowing simple and inexpensive integration with archived samples and existing laboratory workflows. Using human brain metastasis samples, we show that TCR sequences are readily obtained from human tumor tissue and that these sequences are recapitulated by single-cell sequencing methods. This technique will permit detailed studies of the spatial organization of the human T cell repertoire, such as the identification of tumor-infiltrating and tumor-excluded T cell clones.

## Main Text

Spatial transcriptomics permits measurement of gene expression without compromising spatial information. Given the importance of location in many immune processes – such as T cell killing and germinal center formation – the technology promises to transform our understanding of immune function and dysfunction in healthy and diseased tissues. Spatial transcriptomics has been used to describe immune cell infiltrate in diverse tissues including tendons, bone marrow, tumors, joint tissue, and neural tissue^1–6^, and platform commercialization will allow more widespread adoption of the technique^7^. However, detailed understanding of adaptive immune processes requires determination of B and T cell receptor sequences, which determine lymphocyte antigen specificities and can serve as a barcode for lymphocyte clones. Antigen receptor sequencing has transformed single-cell studies of B and T cells^8,9^ but is not yet available in spatial transcriptomics methods. Here, we report a method to determine T cell receptor (TCR) sequences from the commercially available Visium Spatial Gene Expression platform from 10X Genomics. Our technique allows simultaneous visualization of T cell receptor clonotypes and overall gene expression within tissue, uses open-source analysis software, and can be performed on archived cDNA from previous Visium experiments.

The Visium spatial transcriptomics platform uses slide-bound, single-stranded DNA probes to capture polyadenylated mRNA. Each probe contains a partial Read 1 sequence at the 5’ end, a 16-nucleotide spatial barcode, a 12-nucleotide unique molecular identifier (UMI), and a 3’ poly(dT) tail (**Fig, 1a**)^10^. The read 1 primer sequence is initially used for cDNA amplification, the spatial barcode links the probe to a particular spatial location, and the UMI identifies unique cDNA molecules generated during first-strand synthesis. After imaging and tissue permeabilization, a polyadenylated RNA molecule anneals to the poly(dT) tail, allowing subsequent reverse transcription to extend the ssDNA probe to contain the cDNA sequence and an added template switching sequence (**Fig. 1a**). The second strand is generated with a primer complementary to the template switching sequence; this full-length second strand is eluted from the slide by the addition of potassium hydroxide. This eluted second strand is amplified to generate a full-length cDNA library with the spatial barcode and UMI to the 3’ end of the poly(A) tail.

**Figure 1:**
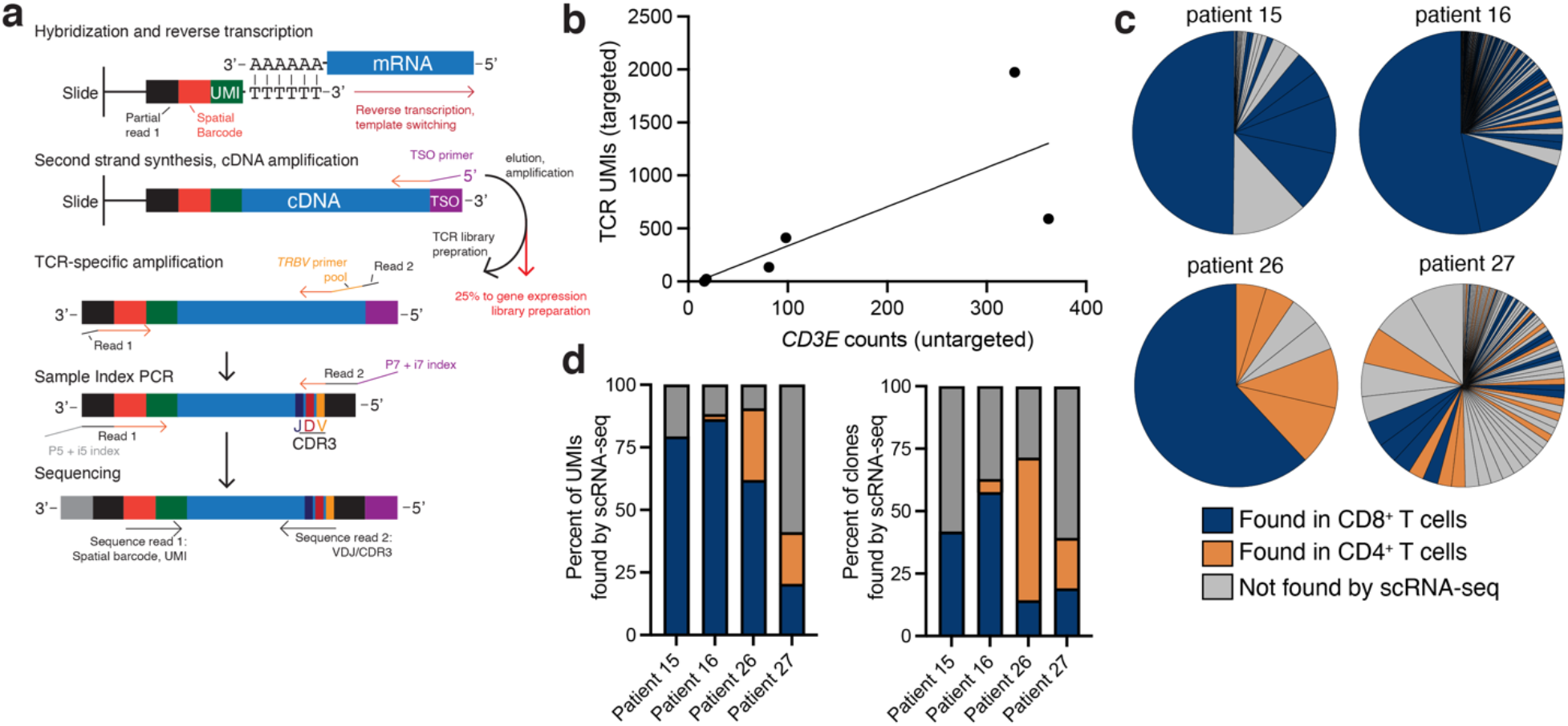
A method for obtaining TCR sequences from the Visium spatial transcriptomics assay. a) Schematic of TCRβ library generation from Visium cDNA. b) Correlation of TCR sequences obtained from the assay with expression of *CD3E* (a pan-T cell marker) by Visium gene expression analysis. c) For four patients, scRNA-seq on T cells was performed to validate TCR sequences obtained by the spatial assay. Pie charts show frequency of clones obtained from the spatial assay and are colored based on whether the clones were independently found through scRNA-seq in CD4^+^ or CD8^+^ T cells. d) Summary of clone recovery statistics. CD4^+^ T cells were not sequenced from patient 15 due to lack of cryopreserved cells.

To generate cDNA molecules containing both the CDR3 region of the TCRβ as well as the UMI and spatial barcodes containing the location information, TCRβ amplification must occur with primers at the 5’ end of the TCR (**Fig. 1a**). Since the TCRβ constant region is 3’ to the CDR3, a pool of variable (*TRBV*) gene primers is required to generate cDNA molecules containing both the CDR3 and spatial information. To generate human libraries, we added a partial Read 2 sequence 5’ to a set of 45 human *TRBV* primers described previously (**Supplementary Table 1**)^11^. PCR was performed with this pool of modified *TRBV* primers and a Read 1 primer using amplified cDNA from the Visium assay as template to generate TCR-enriched libraries. While the standard Visium gene expression protocol calls for fragmentation of the cDNA library^10^, TCR-enriched libraries generated here cannot be fragmented without losing linkage between TCR and spatial barcode sequence. Thus, the TCR-enriched cDNA library was cleaned with bead purification and directly subjected to sample index PCR for multiplexed paired-end sequencing (**Fig. 1a**).

Read 1 of the paired-end sequencing must be performed for at least 28 cycles to provide the sequences of the 16-nucleotide spatial barcode and 12 nucleotide UMI. Read 2 contains the TCRβ-identifying CDR3 sequence and should be preferably longer than 100 nucleotides. In the current study, we performed paired-end sequencing with read lengths ≥150 nucleotides on an Illumina MiSeq instrument. For analysis, read 2 was subjected to the MiXCR pipeline^12^ to identify TCRs present and the reads supporting each identified TCR sequence. UMIs and spatial barcodes were identified from the paired read 1 to correct for amplification bias and to match TCRs to a spatial location.

We performed this protocol and analysis on six fresh-frozen tissue sections of resected human brain metastases, representing four primary tumor types (**Supplementary Table 2**). The number of TCR UMIs identified with this method ranged from 1 to 1,973 per sample, and TCR recovery was loosely correlated both with *CD3E* transcript counts by the standard Visium gene expression assay (**Fig. 1b, Supplementary Table 2**) and CD8^+^ T cell infiltration by flow cytometry (**Supplementary Fig. 1**). The number of unique clones identified ranged from 1 to 113 and diversity (Shannon index) from 0 to 3.89 (**Fig. 1c**). To determine whether the TCRs identified by our technique were recapitulated by other methods, we performed single cell RNA-sequencing (scRNA-seq) on T cells from four matched patients. CD8^+^ T cells were sequenced from all patients, and CD4^+^ T cells were sequenced from three of the four (**Fig. 1c**). On average, 74.8% of TCR UMIs identified by the spatial method described here were also called in matched scRNA-seq samples with the cellranger VDJ pipeline (**Fig. 1d**).

An example implementation of these data is shown in **Figure 2**. A lung cancer metastasis to the brain (patient 27; ref. 13) was stained with hematoxylin and eosin (**Fig. 2a**) and subjected to the standard Visium gene expression assay and spatial TCR method as described above. Gene expression analysis revealed seven clusters of spatial gene expression (**Fig. 2b**). scRNA-seq on FACS-sorted CD4^+^ T cells was performed in parallel, identifying seven clusters of CD4^+^ T cells, including regulatory T cells (T_regs_), granzyme-expressing cells, and naïve cells from a healthy donor (**Fig. 2c-d**). TCR overlap was minimal between these scRNA-seq clusters (**Fig. 2g-h**), and diversity trended higher in CD4^+^ T cells than in CD8^+^ T cells. CD4^+^ T cell clones identified by scRNA-seq were detectable in the Visium TCR results, including single and multiple clones associated with the T_reg_ and granzyme-expressing phenotypes (**Fig. 2e-f**). In Sudmeier et al.^13^, we use this technique to show that TCR clonotypes expressed by exhausted CD8^+^ T cells localize to the tumor parenchyma.

**Figure 2:**
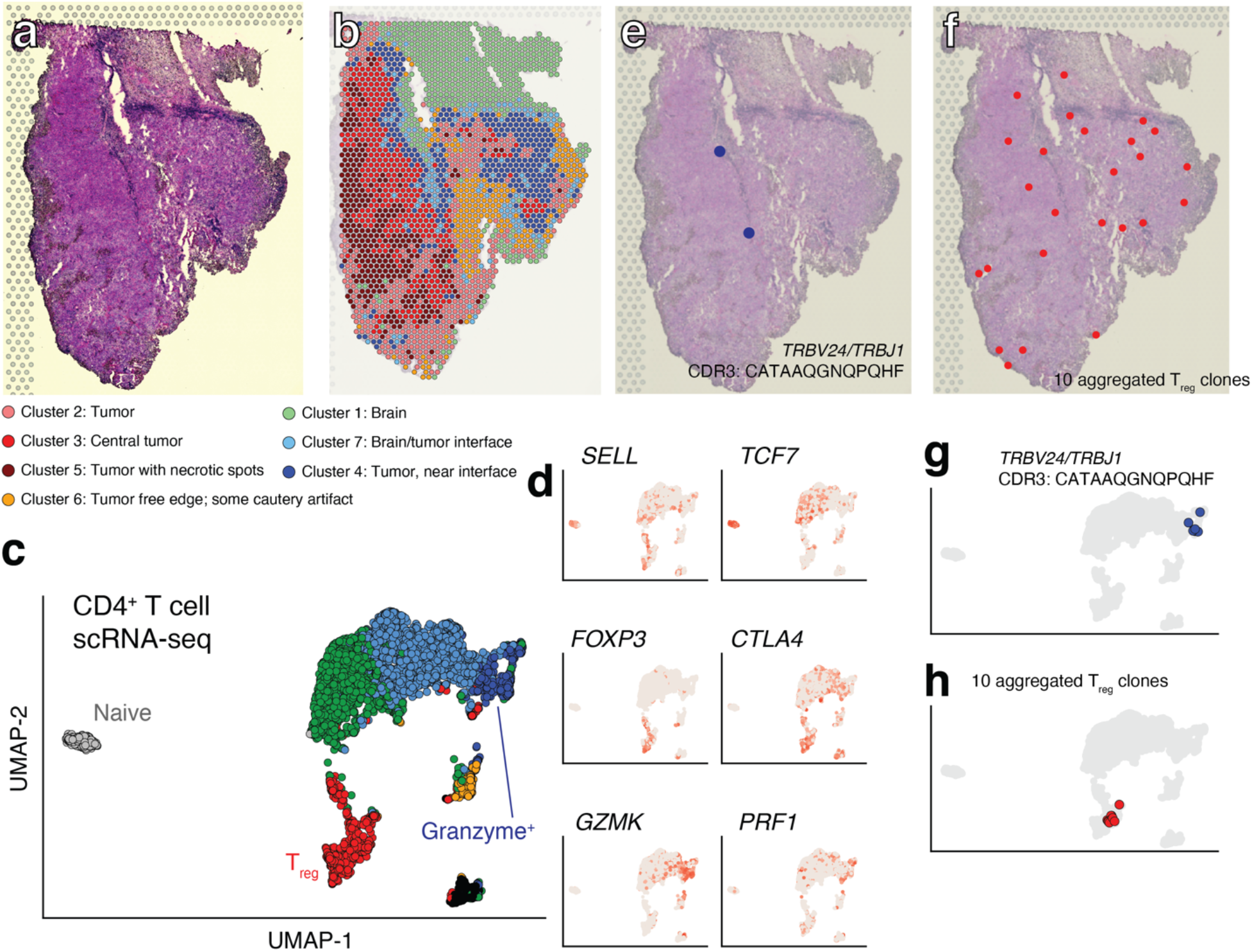
Example spatial localization of CD4^+^ T cell clones. a) H&E staining of a lung cancer metastasis to the brain. b) Clusters of gene expression within the tissue. c) scRNA-seq was performed on CD4^+^ T cells from three patients. Cells are plotted based on their UMAP dimensionality reduction coordinates and colored by gene expression cluster. d) Expression of selected genes. e-f) Spatial localization of a granzyme-expressing CD4^+^ T cell clone and multiple Treg clones. scRNA-seq phenotype of these clones are shown in panels g-h.

Here, we have shown that human TCRβ sequences can be accurately identified and assigned to spatial locations in tumor tissue using the Visium spatial transcriptomics assay. This technique can be used in parallel with the standard gene expression protocol, allowing determination of both TCRβ sequences and the phenotype of the tissue in which they are embedded. The spatial TCR repertoire of tissue types other than cancer can also be explored; for example, the localization of T cell clones in germinal centers or other lymphoid tissues may also be of interest. This method should be generalizable to TCRα sequences by using primer pools specific for *TRAV* genes^11^. Primer pools specific to variable T cell receptor genes of other species should allow this method to be applied to tissue sections from other species (such as mouse^14,15^). Additionally, it should be possible to obtain B cell receptor sequences with this method. Igκ- and Igλ-specific primer pools^16,17^ should generate libraries with insert sizes comparable to the TCR libraries described here, whereas sequencing of heavy chains may require long-read sequencing.

## Acknowledgements

We are grateful to Andreas Wieland for critical reading of the manuscript, to the Emory University School of Medicine Flow Cytometry Core for assistance with cell sorting, to the Yerkes Nonhuman Primate Genomics Core for technical assistance with sequencing, to Jeffrey Olson, Kimberly Hoang, and Edjah Nduom for surgical resection, and to Rafi Ahmed for funding and scientific feedback. W.H.H. is supported by K99AI153736 from the National Institutes of Health (NIH). L.J.S. is supported by a Conquer Cancer Young Investigator Award, a Radiological Society of North America Resident Research Award, and the Nell W. and William Simpson Elkin Fellowship from Emory University. The Yerkes Nonhuman Primate Genomics Core is supported in part by NIH P51OD011132, and Visium gene expression sequencing data was acquired on an Illumina NovaSeq6000 funded by NIH S10OD026799. Research reported in this publication was also supported by the National Cancer Institute of the National Institutes of Health under Award Number P50CA217691. The content is solely the responsibility of the authors and does not necessarily represent the official views of the National Institutes of Health or the American Society of Clinical Oncology^®^ or Conquer Cancer^®^.

## Data deposition

Raw and processed data for scRNA-seq have been deposited in the Gene Expression Omnibus (GEO) under accession GSE179373. Raw and processed data for the gene expression spatial transcriptomics experiments have has been deposited in the GEO with accession GSE179572. Reads from targeted TCR-sequencing for spatial transcriptomics have been deposited in the Sequence Read Archive under accession BioProject PRJNA742564. Sample code for analysis has been posted on Github (https://github.com/whhudson/spatialTCR).

## Methods

### Samples

Detailed information on the patient cohort and acquisition of tumor samples can be found in ref. 13. For spatial transcriptomics, as soon as possible after surgical resection, a piece of tumor tissue was embedded in OCT and flash frozen in a dry ice/2-methybutane bath. 10 μm thick sections were cut and placed onto a 10X Genomics Gene Expression Slide. Slides were stored at −80 °C for up to one week until methanol fixation, H&E staining, and imaging according to the manufacturer’s instructions^18^. Gene expression library preparation was carried out as specified in the manufacturer’s protocol^10^. 5 μl of amplified cDNA from step 3.4 of the manufacturer’s instructions^10^ was used for TCRβ enrichment and library preparation as described in the **Supplementary Protocol**. Data analysis for TCRβ sequencing from Visium is also described in the **Supplementary Protocol**.

Spatial transcriptomics gene expression was performed with 10X Genomics’s spaceranger pipeline. Clusters identified by graph-based clustering were annotated by neuropathologists as described in reference 13.

Remaining tumor not used for spatial transcriptomics was used for cell isolation as described previously^19^: Tumor samples were cut into small pieces in L-15 medium, and digested shaking at 37 °C for 1 hour with a mixture of elastase, DNase, and collagenases I, II, and IV. Samples were forced through a cell strainer and white blood cells were isolated with a 44%/67% Percoll gradient. Cells were counted by flow cytometry and either sorted immediately for scRNA-seq or frozen in FBS with 10% DMSO. For frozen samples, cells were incubated in a 37 °C water bath until thawed and washed twice with RPMI medium supplemented with 10% FBS. For naïve T cell isolation, PBMCs from a healthy donor were used.

For FACS isolation of CD4^+^ and CD8^+^ T cells for scRNA-seq, cells were stained with 1μl of LIVE/DEAD Near IR (Invitrogen L34975), 0.5 tests of anti-CD4 PE/Cy7 (Biolegend 344611), 0.5 tests of anti-CD8 BV421 (Biolegend 344747), 1 test of anti-CD45 FITC (Biolegend 304005), 1 test of anti-CD19 APC (Biolegend 302211), 1 test of anti-CD3 BV786 (Biolegend 344841), and 0.5 tests of anti-CD14 AlexaFluor700 (Biolegend 367113) per million cells. For determination of CD4 and CD8 protein expression, 0.5 tests of anti-CD4 (Biolegend 344651) hashed antibody and 0.5 tests of anti-CD8 hashed antibody (Biolegend 344753) were also used.

For CD8^+^ T cells from tumors, CD45^+^CD14^-^ CD19^-^CD3^+^CD4^-^CD8^+^ cells were isolated by FACS, and for CD4^+^ T cells from tumors, CD45^+^CD14^-^CD19^-^CD3^+^CD4^+^CD8^-^ cells were isolated. CD8^+^ T cells from patients 15 and 16 were sorted at the time of resection. CD4^+^ and CD8^+^ T cells were sorted simultaneously from frozen samples of patients 16, 26, and 27, using hashtag antibodies from Biolegend (product numbers 394661, 394663, 394665, and 394667). TCR data were combined from the frozen and fresh scRNA-seq experiments with patient 16. Naïve CD4^+^ and CD8^+^ T cells from PBMCs were sorted with the strategies above and the addition of CCR7^+^CD45RA^+^ gates (BD 562381 and Biolegend 304134).

### scRNA-seq analysis

The cellranger pipeline (v6.0.0)^20^ was used to align sequencing reads. The Seurat package (v4.0.1) in R was used for data analysis^21^. CD4^+^ T cells were identified by having CITE-seq counts for CD4 > 70 and CITE-seq counts for CD8 < 90. Cells were identified as originating from a particular patient if 90% of its hashtag CITE-seq reads originated from a single barcode. Cells not definitely determined as CD4^+^ or CD8^+^ and/or as from a single patient by CITE-seq were discarded. Doublets were excluded with a maximum cutoff of detected UMIs (10,000), and dead cells were excluded with a maximum percentage of genes originating from mitochondrial genes (5%). TCR genes and Y chromosome genes were excluded from gene expression analysis. Count data were normalized and scaled and variable features identified with Seurat. 10 principal components, 50 neighbors, and a minimum distance of 0 were used for UMAP reduction. Clustering was performed with the Louvain’s algorithm with multilevel refinement and a resolution of 0.3 in Seurat’s FindClusters.

### TCR determination

cellranger vdj was used to call TCR sequences. cellranger vdj gives paired TCRα/TCRβ sequences but only unpaired TCRβ sequences are available from our spatial transcriptomics methods. Thus, we called TCRβ sequences using the same *TRBV* gene family, the same *TRBJ* gene family, and having an identical CDR3 amino acid sequence as an individual clonotype. This method was used for both cellranger vdj and MiXCR output.

### Data visualization

Data were visualized using ggplot2^22^ and GraphPad Prism v9.

### Human subjects

Experiments were carried out with the approval of the Emory University Institutional Review Board under protocols IRB00045732, IRB00095411, and STUDY00001995.

## References

1 Ji, A. L. et al. Multimodal Analysis of Composition and Spatial Architecture in Human Squamous Cell Carcinoma. Cell 182, 497–514.e422, doi:10.1016/j.cell.2020.05.039 (2020).

2 Akbar, M. et al. Single cell and spatial transcriptomics in human tendon disease indicate dysregulated immune homeostasis. Annals of the Rheumatic Diseases, doi:10.1136/annrheumdis-2021-220256 (2021).

3 Baccin, C. et al. Combined single-cell and spatial transcriptomics reveal the molecular, cellular and spatial bone marrow niche organization. Nature Cell Biology 22, 38–48, doi:10.1038/s41556-019-0439-6 (2019).

4 Maynard, K. R. et al. Transcriptome-scale spatial gene expression in the human dorsolateral prefrontal cortex. Nature Neuroscience 24, 425–436, doi:10.1038/s41593-020-00787-0 (2021).

5 Carlberg, K. et al. Exploring inflammatory signatures in arthritic joint biopsies with Spatial Transcriptomics. Scientific Reports 9, doi:10.1038/s41598-019-55441-y (2019).

6 Ståhl, P. L. et al. Visualization and analysis of gene expression in tissue sections by spatial transcriptomics. Science 353, 78–82, doi:10.1126/science.aaf2403 (2016).

7 Marx, V. Method of the Year: spatially resolved transcriptomics. Nature Methods 18, 9–14, doi:10.1038/s41592-020-01033-y (2021).

8 De Simone, M., Rossetti, G. & Pagani, M. Single Cell T Cell Receptor Sequencing: Techniques and Future Challenges. Frontiers in Immunology 9, doi:10.3389/fimmu.2018.01638 (2018).

9 Papalexi, E. & Satija, R. Single-cell RNA sequencing to explore immune cell heterogeneity. Nature Reviews Immunology 18, 35–45, doi:10.1038/nri.2017.76 (2017).

10 Genomics, X. VisiumSpatial Gene Expression Reagent Kits, <https://assets.ctfassets.net/an68im79xiti/3GGIfH3RWpd1bFVha1pexR/8baa08d9007157592b65b2cdc7130990/CG000239_VisiumSpatialGeneExpression_UserGuide_RevD.pdf>(

11 Robins, H. S. et al. Comprehensive assessment of T-cell receptor β-chain diversity in αβ T cells. Blood 114, 4099–4107, doi:10.1182/blood-2009-04-217604 (2009).

12 Bolotin, D. A. et al. MiXCR: software for comprehensive adaptive immunity profiling. Nature Methods 12, 380–381, doi:10.1038/nmeth.3364 (2015).

13 Sudmeier, L. J. et al. The CD8+ T cell landscape of human brain metastases. (Submitted).

14 Saligrama, N. et al. Opposing T cell responses in experimental autoimmune encephalomyelitis. Nature 572, 481–487, doi:10.1038/s41586-019-1467-x (2019).

15 Dash, P. et al. Paired analysis of TCRα and TCRβ chains at the single-cell level in mice. Journal of Clinical Investigation 121, 288–295, doi:10.1172/jci44752 (2011).

16 Tiller, T. et al. Efficient generation of monoclonal antibodies from single human B cells by single cell RT-PCR and expression vector cloning. Journal of Immunological Methods 329, 112–124, doi:10.1016/j.jim.2007.09.017 (2008).

17 von Boehmer, L. et al. Sequencing and cloning of antigen-specific antibodies from mouse memory B cells. Nature Protocols 11, 1908–1923, doi:10.1038/nprot.2016.102 (2016).

18 Genomics, X. Methanol Fixation, H&E Staining & Imaging for Visium Spatial Protocols, <https://assets.ctfassets.net/an68im79xiti/2r4GCKmP3nHxaGnYYr46Di/fe3215affa772282dcf486932de7ac84/CG000160_DemonstratedProtocol_MethanolFixationandHEStaining_RevB.pdf> (

19 Wieland, A. et al. Defining HPV-specific B cell responses in patients with head and neck cancer. Nature, doi:10.1038/s41586-020-2931-3 (2020).

20 Zheng, G. X. Y. et al. Massively parallel digital transcriptional profiling of single cells. Nature Communications 8, doi:10.1038/ncomms14049 (2017).

21 Hao, Y. et al. Integrated analysis of multimodal single-cell data. Cell 184, 3573–3587.e3529, doi:10.1016/j.cell.2021.04.048 (2021).

22 Wickham, H. Ggplot2: elegant graphics for data analysis. (Springer, 2009).

