## Supplementary Information for "Localization of T cell clonotypes using spatial transcriptomics"

**Supplementary Table 1.** See attached Excel file. 45 *TRBV* primers with partial read 2 sequences at the 5' end. The partial read 1 sequence for cDNA amplification is also given.

| | Sample 15 | Sample 16 | Sample 19 | Sample 24 | Sample 26 | Sample 27 |
|---|---|---|---|---|---|---|
| # reads | 509157* | 2057866 | 348486 | 754089 | 1130748 | 2993934 |
| #TCR UMIs | 590 | 1973 | 1 | 135 | 21 | 411 |
| # clones | 24 | 113 | 1 | 44 | 7 | 84 |
| TCR UMIs/spot | 0.28 | 1.15 | 0.001 | 0.10 | 0.005 | 0.19 |
| *CD3E* counts (Visium; untargeted) | 362 | 328 | 16 | 81 | 18 | 98 |
| TCR sequencing saturation | 98.4% | 98.1% | >99.9% | 98.7% | >99.9% | >99.9% |
| $CD8^+$ cells/g tumor ($x10^5$) | 3.76 | 23.07 | 0.87 | 0.96 | 1.78 | 30.38 |
| Primary tumor type | Lung | Melanoma | Lung | Renal clear cell | Breast | Lung |

*Sample was sequenced on two separate MiSeq runs.

**Supplementary Table 2.** Statistics for the six samples on which spatial TCR sequencing was performed.

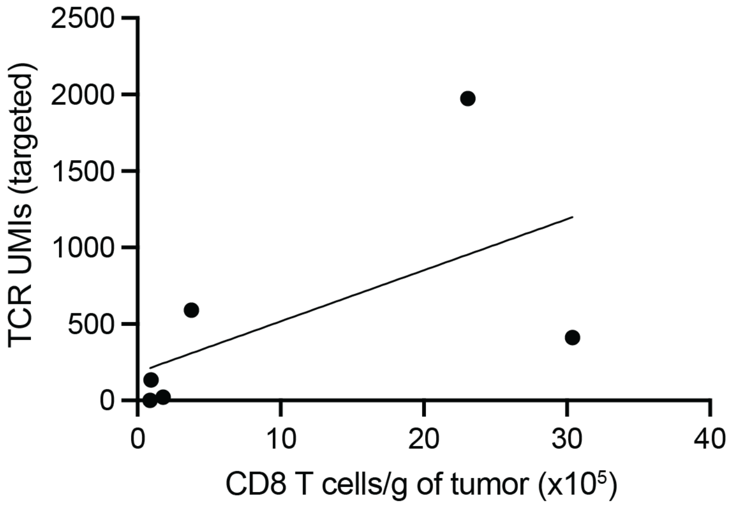

**Supplementary Figure 1.** Correlation of TCR UMIs obtained from targeted spatial TCR sequencing with $CD8^+$ T cell infiltration by flow cytometry.

**Supplementary Protocol**

TCR library preparation

**Materials**

1. Nuclease free-water (Ambion AM9932)
2. 45 human *TRBV* specific primers with 5' partial read 2 sequences (see **Table** for sequences), resuspended in Nuclease-free water. We ordered these primers separately from Integrated DNA Technologies in a 96-well plate format with the complete yield resuspended to 100 μM.
3. Amplified cDNA from step 3.4 of the Visium Spatial Gene Expression Reagent Kits User Guide[1].
4. 10X Genomics Library Construction kit (PN-1000190)
5. PCR tubes, DNase/RNAse-free
6. Thermocycler
7. SPRIselect beads (Beckman Coulter B23317)
8. 80% ethanol prepared with nuclease-free water
9. Buffer EB (Qiagen catalog 19086)
10. (Optional) Qubit instrument with dsDNA HS Assay kit
11. 10X Genomics Dual Index Kit TT Set A sample index primers (PN-1000215)

**Method**

This protocol begins with amplified cDNA resulting from step 3.4 from the Visium Spatial Gene Expression Reagent Kits User Guide[1]. 10 μl of amplified cDNA is used for gene expression library preparation, and we here use 5 μl of amplified cDNA for amplification and preparation of TCR libraries. This amount may be adjusted to conserve cDNA or provide greater sequencing depth. While 10X Genomics recommends storage of amplified cDNA at -20 °C for up to four weeks, we have successfully prepared TCR spatial libraries from amplified cDNA frozen for longer periods.

1. Generate the pool of *TRBV* primers by combining 45 *TRBV*-specific primers (see **Supplementary Table 1**) in equimolar amounts. If ordered as in point (2) of Materials, equal volumes of each primer can be added.
2. Dilute primer pool to a concentration of 10 μM total concentration of primer (0.22 μM of each primer). If prepared as step (1), add nuclease-free water at a 9:1 ratio to the pooled primers generated in step (1). Also dilute partial read 1 primer (**Supplementary Table 1**) to 10 μM.
3. Prepare TCR cDNA amplification mix in a PCR tube as outlined below. Scale linearly for different amounts of amplified cDNA input.

| | |
|---|---|
| Amplification mix (from 10X Genomics Library Construction Kit) | 7.14 µl |
| Amplified cDNA | 5 µl |
| *TRBV* primer pool (10 µM total) | 0.29 µl |
| Partial read 1 primer (10 µM) | 0.29 µl |
| Nuclease-free water | 1.57 µl |
| Total | 14.29 µl |

4. Amplify TCR cDNA with the following cycling conditions, repeating steps 2-4 for 35 cycles. Cycle number may be adjusted downward for large amounts of cDNA. The KAPA SYBR FAST qPCR kit may be used to determine amplification in real-time, if desired.

| | | |
|---|---|---|
| 1. Initial denaturation | 98 °C | 03:00 (mm:ss) |
| 2. Denaturation | 98 °C | 00:15 |
| 3. Annealing | 59 °C | 00:20 |
| 4. Extension | 72 °C | 01:00 |
| 5. Final extension | 72 °C | 01:00 |
| 6. Hold | 4 °C | ∞ |

5. Perform cleanup of PCR product with SPRIselect beads as follows:
   a. (Optional) Bring PCR reaction volume up to 50 µl with nuclease-free water (by adding 35.7 µl if prepared as in step (3)).
   b. Ensure that SPRIselect beads are well mixed by shaking. Add 0.6X of SPRIselect beads to PCR reaction; 30 µl if using a 50 µl volume as above.
   c. Mix well by pipetting.
   d. Incubate 5 minutes at room temperature.
   e. Place on separation magnet until solution clears.
   f. Pipette to remove and discard supernatant.
   g. Add 200 µl of 80% ethanol to the pellet. Do not remove tube from magnet or pipette to mix. Wait 30 seconds.
   h. Pipette to remove and discard supernatant.
   i. Repeat steps g-h for a total of two 80% ethanol wash steps.
   j. Ensure that all 80% ethanol is removed. Air dry for two minutes.
   k. Remove tube from magnet.
   l. Add 40.5 µl of Buffer EB to tube. Resuspend beads by pipette mixing.
   m. Incubate 2 minutes at room temperature.
   n. Place on separation magnet until solution clears.
   o. Transfer 40 µl to a new PCR tube. This is the amplified TCR cDNA.

6. Store amplified TCR cDNA at 4 °C overnight, freeze at -20 °C for longer storage, or continue to next step.
7. Recommended: quantify amplified TCR cDNA using a Qubit High Sensitivity Assay kit according to manufacturer's instructions. Our yields ranged from 50-150 ng of total product.
8. Prepare sample indexing PCR reaction for library generation and multiplexed sequencing in a PCR tube as outlined below:

| | |
|---|---|
| TCR cDNA from step 6 | 15 µl |
| TT set A primers | 10 µl |
| Amplification mix (from 10X Genomics Library Construction Kit) | 25 µl |
| Total | 50 µl |

9. Amplify TCR cDNA with the following cycling conditions, repeating steps 2-4 for 15 cycles:

| | | |
|---|---|---|
| 1. Initial denaturation | 98 °C | 00:45 (mm:ss) |
| 2. Denaturation | 98 °C | 00:20 |
| 3. Annealing | 67 °C | 00:30 |
| 4. Extension | 72 °C | 01:00 |
| 5. Final extension | 72 °C | 01:00 |
| 6. Hold | 4 °C | Hold |

Note: cycle number for steps 2-4 may be optimized based on yield from step 7.

10. Perform cleanup of PCR product with SPRIselect beads as in step 5.

Sequencing

We performed sequencing with the Yerkes Nonhuman Primate Genomics Core at Emory University. Libraries were validated by capillary electrophoresis on a Bioanalyzer (Agilent), pooled at equimolar concentrations, and sequenced on an Illumina MiSeq using a MiSeq Reagent Kit v3 at PE300 with 10% PhiX.

Sequence Analysis

Paired-end sequencing will result in the generation of two FASTQ files. Read 1 contains the spatial barcode and UMI and read 2 contains the VDJ sequences. We used MiXCR[2] to call antigen receptor sequences from read 2 and used a custom Python 2 script to perform UMI correction and count number of antigen receptor UMIs at a particular spatial location.

**Files needed:**

Visium tissue positions list. This is typically found in Space Ranger output as spatial/tissue_positions_list.csv after running the Space Ranger gene expression pipeline.

Read 1 and 2 FASTQ files from sequencing section, above.

We have provided example downsampled FASTQ files, a Visium tissue positions list, a sample python script (combine_counts.py), and expected output from this pipeline on Github (https://github.com/whhudson/spatialTCR).

**Software needed:**

1. MiXCR (version 3.0.13 is the latest release as of writing)
2. (Optional) Python

**Method:**

1. Use the mixcr analyze pipeline to call antigen receptor sequences from the read 2 FASTQ file. The following code is provided as an example. path/to/read2.fastq is the file path to the read 2 FASTQ file above. The saveOriginalReads parameter should be set to true and alignments should be written to allow for extraction of sequencing reads in the next step.

```
mixcr analyze shotgun -s hsa --starting-material rna --align "-OsaveOriginalReads=true" --assemble "--write-alignments" path/to/read2.fastq.gz mixcr_output/clones
```

2. Export individual reads supporting each clone with the mixcr exportReadsForClones option. The following code is provided as an example. This will result in fastq.gz files with reads supporting each clone being written into the mixcr_output/reads folder as reads_clnN.fastq.gz, where N is the clone number identifier.

```
mixcr exportReadsForClones -s mixcr_output/clones.clns mixcr_output/reads/reads.fastq.gz
```

   At this point, clones have been identified and mapped to individual sequencing reads. Information about each clone will be given in the mixcr_output/clones.clonotypes.ALL.txt file if the naming convention above has been followed. Additionally, the sequencing read 2 for each clone has been written into the mixcr_output/reads folder. At this point, the UMI and spatial barcode from the paired read 1 should be extracted to map each TCR read to a UMI and spatial location. We provide the example Python 2 script combine_counts.py for performing this task.

3. In the combine_counts.py script, specify the directory containing the reads written by mixcr (path/to/mixcr_output/reads if using the naming as in step 2 above), the tissue positions list from 10X Genomics Space Ranger output, and the path to the read 1 FASTQ from the sequencing step above.

4. Running the script will output a tab-delimited text file with each spot/spatial barcode as a row and each clone as a column. The number in each cell indicates the number of UMIs measured in each spatial barcode (row) for a particular clone (column). This tab-delimited text file can then be used for further analysis in R, such as to add metadata to a Seurat spatial object.

**References**


1 Genomics, X. *VisiumSpatial Gene Expression Reagent Kits*, <https://assets.ctfassets.net/an68im79xiti/3GGIfH3RWpd1bFVha1pexR/8baa08d9007157592b65b2cdc7130990/CG000239_VisiumSpatialGeneExpression_UserGuide_RevD.pdf> (

2 Bolotin, D. A. *et al.* MiXCR: software for comprehensive adaptive immunity profiling. *Nature Methods* **12**, 380-381, doi:10.1038/nmeth.3364 (2015).
